# A novel single-color FRET biosensor for Rho-kinase activity reveals calcium-dependent activation of RhoA and ROCK

**DOI:** 10.1101/2024.05.30.596680

**Authors:** Allison E. Mancini, Megan A. Rizzo

## Abstract

Ras homolog family member A (RhoA) acts as a signaling hub in many cellular processes, including cytoskeletal dynamics, division, migration, and adhesion. RhoA activity is tightly spatiotemporally controlled, but whether downstream effectors share these activation dynamics is unknown. We developed a novel single-color FRET biosensor to measure Rho-associated kinase (ROCK) activity with high spatiotemporal resolution in live cells. We report the validation of the Rho-Kinase Activity Reporter (RhoKAR) biosensor. RhoKAR activation was specific to ROCK activity and was insensitive to other kinases. We then assessed the mechanisms of ROCK activation in mouse fibroblasts. Increasing intracellular calcium with ionomycin increased RhoKAR activity, and depleting intracellular calcium with EGTA decreased RhoKAR activity. We also investigated the signaling intermediates in this process. Blocking calmodulin or CaMKII prevented calcium-dependent activation of ROCK. These results indicate that ROCK activity is increased by calcium in fibroblasts and that this activation occurs downstream of CaM/CaMKII.

## INTRODUCTION

Ras homolog family member A (RhoA) is a small GTPase that acts as a molecular switch and signaling hub, controlling integral processes including migration, cell cycle progression, and cell division^1–5^. Like other small GTPases, RhoA cycles between an active, GTP-bound “on” state, localized to the membrane, and an inactive, GDP-bound “off” state, localized to the cytosol ^6–10^. This cycling is controlled mainly by activating RhoGEFs that promote the exchange of GDP for GTP and inactivating RhoGAPs that enhance GTP hydrolysis^6,9,11–17^. Additionally, RhoGDIs bind and sequester inactive RhoA in the cytoplasm and extract RhoA from the membrane following GTP hydrolysis^6,18–22^. Together, these proteins respond to diverse signaling events to finely tune RhoA activity.

RhoA’s activity is mediated by several downstream effectors, with its major effector being Rho-associated kinase (ROCK). ROCK is a serine-threonine protein kinase that can phosphorylate its targets when bound to active RhoA. ROCK’s targets include adducin, ERM family members, LIM kinase, myosin light chain phosphatase, the Na^+^/H^+^ exchanger, and vimentin, among others^23–25^. ROCK has emerged in recent years as a potential therapeutic target in diseases including metastatic cancer, stroke, Alzheimer’s disease, and spinal cord injury^26–28^. Clinically, Rho-kinase inhibitors manage ophthalmologic conditions, including glaucoma, ocular hypertension, and diabetic retinopathy^29,30^. ROCK inhibitors have also been successfully used to manage vascular conditions such as erectile dysfunction and pulmonary hypertension, as well as to improve chronic graft-versus-host disease^31–37^.

Förster resonance energy transfer (FRET) biosensors sensitively quantify protein activity in living cells, revealing the spatial and temporal dynamics of key molecular signals^38,39^. Typically, FRET sensors incorporate donor and acceptor fluorophores of different colors, notably cyan and yellow fluorescent proteins^39,40^. Changes in sensor activity alter the FRET efficiency, which is typically quantified by using the acceptor-to-donor fluorescence ratio. The direct relationship between FRET efficiency and sensor activity allows calibration and quantitative analyses^41–43^. Even so, a two-color, heterotransfer approach to FRET has several drawbacks, most notably the large portion of the visible spectrum taken up by using two differently colored fluorophores. These reporters are challenging to combine with other optical tools, such as channelrhodopsin, additional sensors, and organic fluorescent indicator dyes.

To improve quantification and multiplexing, we developed single-color FRET biosensors that measure a change in the polarization state rather than the color of the emitted light. These FLuorescence Anisotropy REporters (FLAREs) take advantage of the inherent polarized emission from all fluorescent proteins and measure FRET as a decrease in fluorescence anisotropy^44–46^. Because fluorescence anisotropy can be measured more sensitively than fluorescence intensity and because these sensors are well suited for multiplexing, these homotransfer reporters are especially well-suited for applications where the activity of two or more proteins need to be quantitated precisely in the same cell at the same time^47^. Further, restriction to a single color creates space in the visible spectrum for optogenetic tools, furthering the possibilities for cellular manipulation during imaging experiments.

Because of the increasing interest in ROCK’s role in driving clinical pathologies and basic cell physiology, we developed a sensor that reports ROCK activity in living cells. Although two-color FRET-based biosensors exist for several members in the RhoA signaling cascade, including RhoA, RhoGEFs, RhoGAPs, RhoGDI, and ROCK, the two-color approach limits possibilities for multiplexing sensors to study the activity of multiple proteins in real-time ^48–53^. Furthermore, a previous ROCK FRET sensor showed cross-reactivity to PKA, hampering its utility in studying ROCK specifically^53^. To investigate the role of ROCK in live cells in a spatiotemporally resolved manner using a polarization-based FRET approach, we have developed a single-color ROCK FLARE-type biosensor, the Rho-Kinase Activity Reporter (RhoKAR). The sensor gives a readout specific to ROCK activity and is insensitive to the activity of other kinases. This sensor has also revealed calcium-dependent activation of ROCK, underscoring the utility of this tool in characterizing signaling behavior in live cells.

## METHODS

| Reagent Type or Resource | Designation | Source or Reference | Identifier |
| --- | --- | --- | --- |
| Cell line | NIH-3T3 |  | RRID: CVCL_0594 |
| Chemical compound, drug | Lysophosphatidic acid | Sigma-Aldrich | L7260 |
| Chemical compound, drug | Y27632 | Sigma-Aldrich | Y0503 |
| Chemical compound, drug | Forskolin | Sigma-Aldrich | F6886 |
| Chemical compound, drug | Ionomycin calcium salt from *Streptomyces conglobatus* | Sigma-Aldrich | I0634 |
| Chemical compound, drug | EGTA | Research Projects International | E14100 |
| Chemical compound, drug | H89 | Cell Signaling Technologies | 9844 |
| Strain (*Escheria coli*) | JM109 Competent Cells | Promega | L2005 |
| Commercial kit | Miniprep Kit | Qiagen | 27104 |
| Commercial kit | HiSpeed Midiprep Kit | Qiagen | 12643 |
| Software | Zeiss AxioVision |  | RRID: SCR_002677 |
| Software | Zeiss ZEN Blue |  | RRID: SCR_013672 |
| Software | FIJI |  | RRID: SCR_002285 |
| Software | Microsoft Excel |  | RRID: SCR_016137 |
| Software | GraphPad Prism |  | RRID: SCR_002798 |
| Software | BioRender |  | RRID: SCR_018361 |

### Plasmid and construct cloning

The RhoKAR sensor was generated using standard molecular cloning techniques. Briefly, an insert containing an FHA domain and the ROCK substrate from adducin was ordered as an oligo inserted in the pUC57-Kan backbone from GenScript with XhoI and AgeI cut sites flanking the insert. The plasmid was transformed into JM109 competent *E. coli* via standard heat shock technique. These cells were then grown in media containing kanamycin to select for cells carrying the plasmid. The plasmid was extracted using a standard DNA isolation Miniprep kit (Qiagen). Using XhoI and AgeI restriction enzymes on the isolated plasmid, the insert of interest was excised via enzymatic digest, gel purified, and then ligated into the tandem mVenus:mVenus FRET backbone previously developed by our lab^54^. The presence of the insert in the correct location and orientation within the mVenus:mVenus tandem FRET backbone was confirmed with a test digest and visualization of an insert of the appropriate size via gel electrophoresis, as well as via Sanger sequencing.

### Cell culture and transfection

NIH-3T3 cells were maintained in Dulbecco’s Modification of Eagle’s Medium (DMEM) supplemented with 10% fetal bovine serum and 1% penicillin/streptomycin. Cells were plated on 35-mm glass bottom dishes and incubated at 37°C with 5% CO_2_ 24 hours before transfection. Cells were transfected using 1 µg of DNA per plate using LipoD293 (SignaGen) 48 hours before imaging. The growth media was removed immediately before imaging, and cells were washed with Hank’s Balanced Salt Solution (HBSS) with 0.1% bovine serum albumin (BSA). Cells were maintained in HBSS with 0.1% BSA for short time course imaging experiments. Where indicated, serum starvation was achieved by replacing the complete media with serum-free DMEM overnight before imaging.

### Fluorescence polarization microscopy

Fluorescence polarization methods are described in detail in previous publications^47,55^. Briefly, two Zeiss AxioObserver Microscopes (Carl Zeiss MicroImaging) equipped for widefield imaging were used to gather fluorescence anisotropy images. Images were collected with a 20X 0.75 NA objective lens on one microscope. This setup included a 505 nm LED illumination source, followed by a wire grid polarizer (Meadowlark Optics) in the excitation pathway before a filter turret containing a YFP filter cube (Zeiss). Light from the sample passes through a DualView beam splitter (Optical Insights) equipped with filters for the parallel and perpendicular states of light before being collected as a single image by an Orca-R2 water-cooled camera (Hamamatsu). For CFP-YFP FRET experiments, this same microscope setup was used, with a 455 nm LED illumination source, a CFP-YFP filter cube in the turret, and a CFP/YFP filter set in the DualView. The second Zeiss AxioObserver had a nearly identical setup but used a Gemini W-View (Hamamatsu) as a beam splitter before light collection with an AxioCam 506 (Zeiss) camera. Where indicated, a 40X 1.3 NA oil immersion objective lens was used for imaging on the second microscope.

### Image analysis

Widefield images of different polarization states were split into separate images, then recombined into a single two-channel image and aligned using the StackReg plugin package in FIJI^56^. Background-corrected fluorescence intensity measurements were taken at each time point in individual cells using hand-drawn ROIs. Fluorescence anisotropies were calculated as described previously. Briefly, fluorescence anisotropy (R) is calculated as a ratio between the intensities of the two polarization states of light, parallel (P) and perpendicular (S), over the total fluorescence intensity:

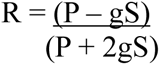

where (g) corrects for differences in camera sensitivity to each polarization state of light. After collecting fluorescence intensity measurements in FIJI, fluorescence anisotropies were calculated in Microsoft Excel, and statistical analysis and figure generation were done using GraphPad Prism. Because a decrease in anisotropy (R) corresponds to an increase in ROCK activity for the RhoKAR sensor, fluorescence anisotropies are presented as -R in this paper. For LUT image creation, background-corrected fluorescence anisotropies were calculated in each pixel within FIJI using the above equation, multiplied by −1 to get -R, backgrounds were removed, images were smoothed, and the MPL-Inferno LUT was applied to create FRET map images. Figures were compiled in BioRender and Microsoft PowerPoint.

## RESULTS

### Development and validation of single-color RhoKAR FRET biosensor

FLAREs use two fluorophores of a single color to provide homotransfer-FRET readouts based on the change in the polarization state of emitted light. Fluorescence emission from fluorophores is highly polarized or anisotropic^44^. When light in a uniform polarization state excites a fluorophore, only fluorophores in the correct geometric orientation in space are excited via a process known as photoselection (**Figure 1A, 1B**)^46^. Emission from photoselected fluorophores will be in the same polarization state as the exciting wavelength without energy transfer to a nearby acceptor fluorophore. However, acceptor emission is depolarized if energy transfer occurs to an acceptor fluorophore within a molecular distance (**Figure 1C**). This technique permits the design of single-color FRET biosensors that provide real-time protein activity readouts in live cells. Fluorescence anisotropies can also be measured more sensitively than fluorescence intensity^44^, allowing precise quantification of even small shifts in FRET ratios and, thus, protein activity. FLAREs do not require bleed-through correction and can be more easily multiplexed than heterotransfer FRET sensors^46^. FLAREs can be used on standard widefield microscopes using a linear polarizer in the light path before excitation of the sample and a polarization-splitting optical filter in front of the camera. This setup allows for the collection of light in both polarization states simultaneously to get fluorescence anisotropy readouts of biosensor activity^48,55^.

**Figure 1:**
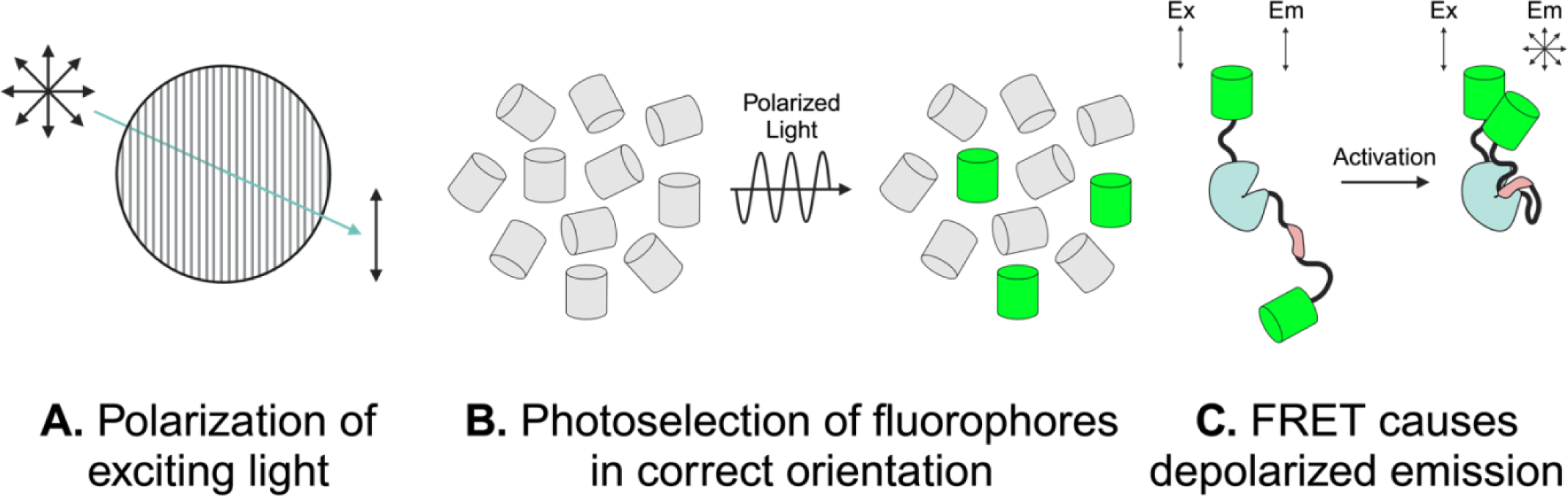
Anisotropy-based FRET strategy. **A**: Light passes through a polarized filter, so all exciting light is in the same polarization state. **B**: The polarized light excites fluorophores in the correct geometric orientation. **C**: In the absence of FRET, emission from photoselected fluorophores will be in the same polarization state as exciting light. Upon conformational change of the biosensor associated with activation, FRET to a nearby acceptor fluorophore causes depolarized emission. Adapted from Snell *et al*.^46^

We developed a single-color FRET sensor, the RhoKAR sensor, to measure ROCK activity directly. RhoKAR consists of two mVenus molecules flanking an FHA domain and the phosphorylation consensus sequence from adducin, a target of ROCK (**Figure 2A, 2B**). Activated ROCK phosphorylates the sensor’s substrate, resulting in binding to the adjacent FHA domain and a conformational change that increases FRET. Images of the sensor were collected in the perpendicular (*S*) and parallel (*P*) polarization states (**Figure 2C**). We then calculated fluorescence anisotropies (*-R*) using the difference in intensities in the two channels. We visualized RhoKAR activity throughout the cell with a pseudo-colored FRET map image generated in FIJI.

**Figure 2:**
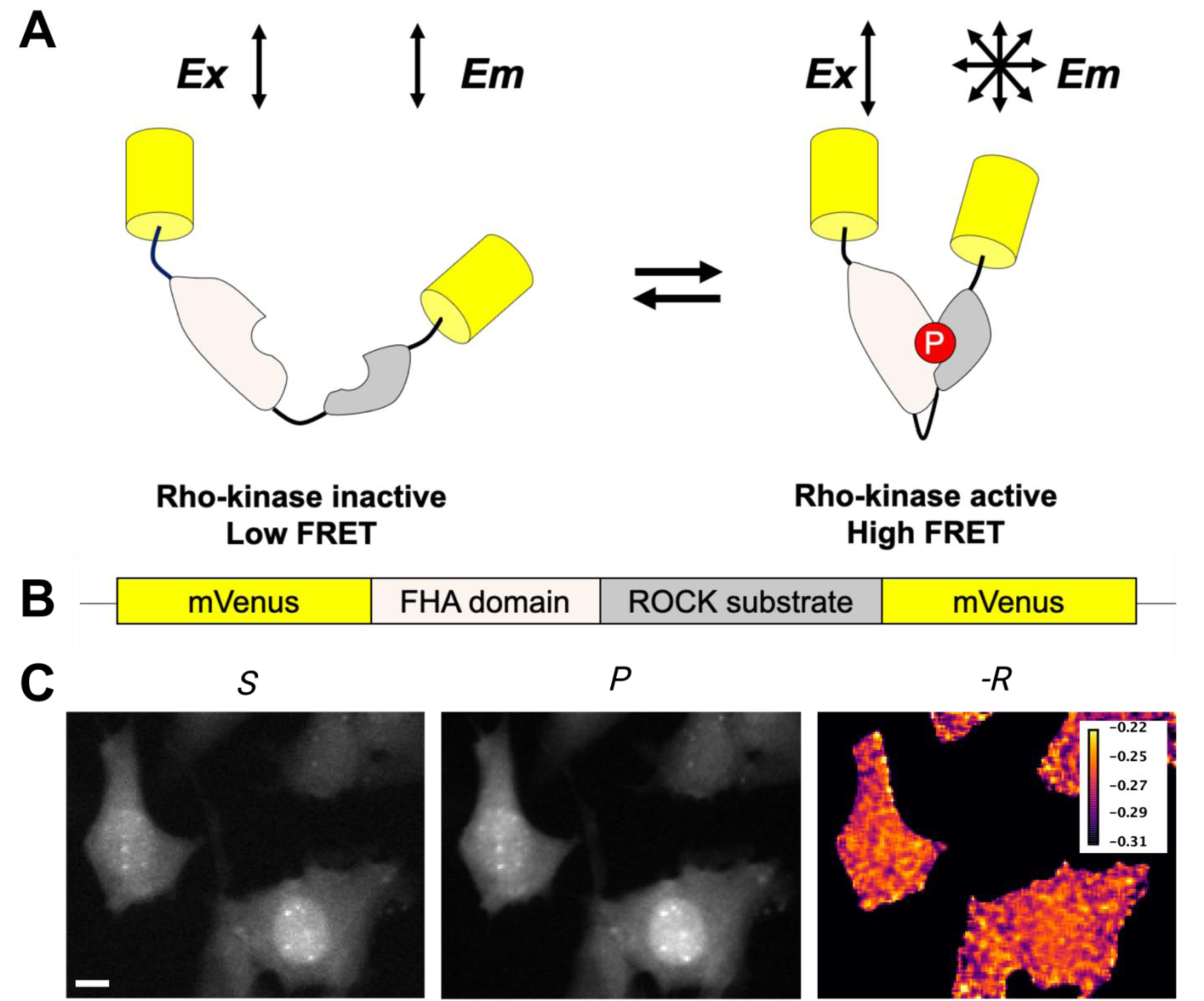
RhoKAR sensor schematic. **A**: Active ROCK phosphorylates the substrate, leading to a conformational change and an increase in FRET. **B**: The RhoKAR sensor consists of two mVenus domains flanking an FHA domain and the ROCK substrate from adducin. **C**: Representative images of NIH-3T3 cells expressing the RhoKAR sensor in the Perpendicular (*S*) and Parallel (*P*) polarization state channels and pseudo-colored FRET map (*-R*), where brighter color indicates higher ROCK activity. Scale bar = 10 µm.

### RhoKAR provides a readout of ROCK stimulation and inhibition

To determine whether the sensor responds appropriately to changes in ROCK activity, we stimulated serum-starved cells expressing RhoKAR with 6 µM lysophosphatidic acid (LPA), a classical RhoA activator^57^, during a short imaging time course. We imaged cells for several minutes before drug addition to establish baseline anisotropies and added the indicated drug at T = 0 minutes. The average fluorescence anisotropy in each cell in the 180 seconds before drug addition was subtracted from the fluorescence anisotropy in that same cell at each time point to yield a change in fluorescence anisotropy (−ΔR) (**Figure 3A**). LPA treatment significantly increased the change in fluorescence anisotropy over baseline in the RhoKAR sensor compared to equivalent volume dH_2_O vehicle control, corresponding to an increase ROCK activity and phosphorylation of the adducin substrate on RhoKAR (**Figure 3B**). Intensity maps with FRET map overlays show increased ROCK activity at 2 minutes compared to the time point immediately before LPA addition (**Figure 2C**). Next, we treated cells with 5 µM Y-27632, a selective ATP-competitive inhibitor of ROCK (**Figure 3D**). Y27632 treatment significantly decreased RhoKAR activity compared to vehicle control (**Figure 3E**). FRET map images illustrate the decrease in RhoKAR activity seen with the Y27632 treatment (**Figure 3F**). Because the measured anisotropy shifts as expected when ROCK is stimulated or inhibited, these results show that the RhoKAR sensor reveals both ROCK activation and de-activation in live cells.

**Figure 3:**
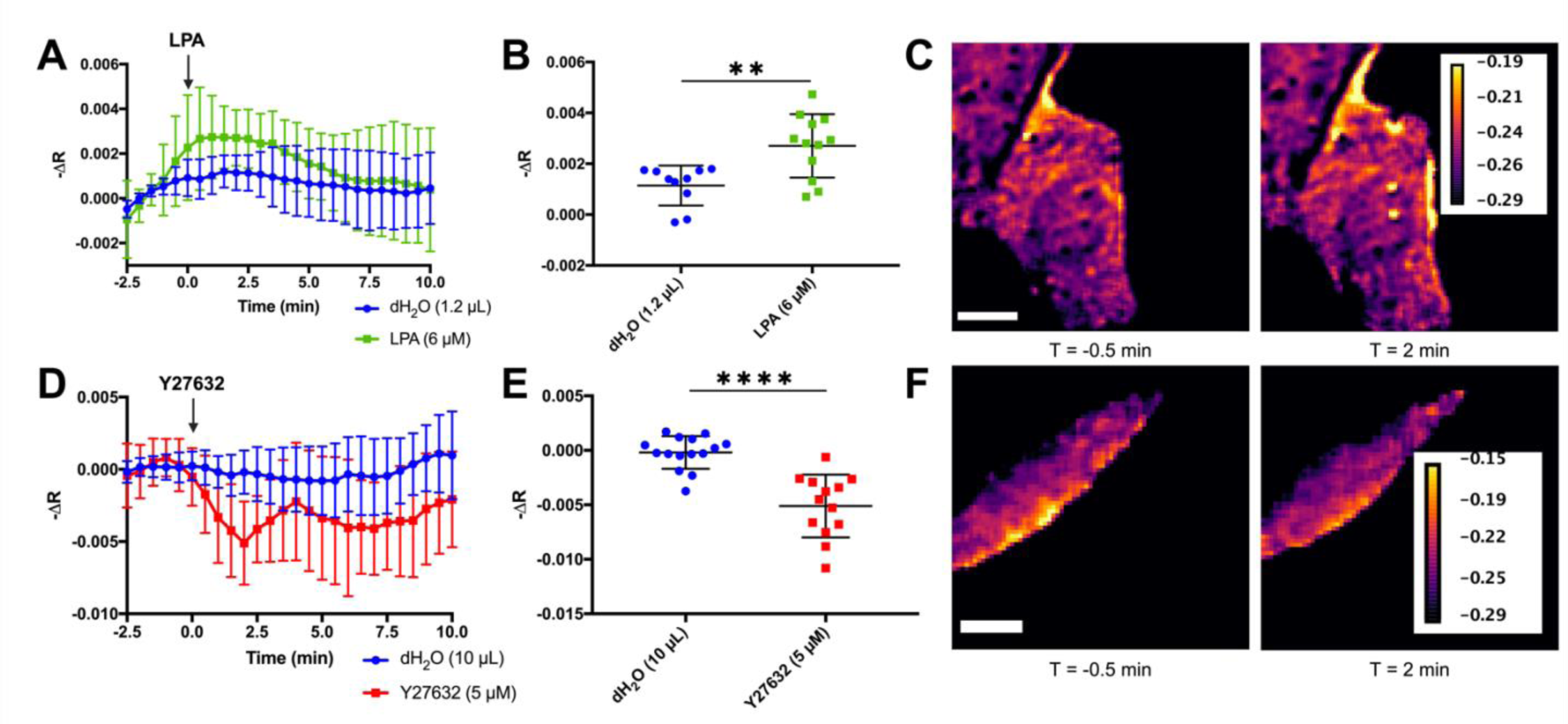
The RhoKAR sensor gives readout of ROCK stimulation and inhibition. **A**, **B:** Time-course change in fluorescence anisotropies in individual NIH-3T3 cells expressing the RhoKAR sensor when stimulated with 6 µM LPA or 1.2 µL dH_2_O vehicle control (t-test, **p<0.01, n = 10 for dH_2_O, n = 12 for LPA). **C**: Representative FRET map image of a cell immediately before and 5 minutes after LPA treatment. Scale bar = 10 µm. **D**, **E**: Time-course change in fluorescence anisotropies in individual cells expressing the RhoKAR sensor when stimulated with Y27632 or dH_2_O vehicle control (t-test, ****p<0.0001, n = 15 for dH_2_O, n = 13 for Y27632). **F**: Representative FRET map of cells immediately before and 2 minutes after Y27632 treatment. Scale bar = 10 µm.

### RhoKAR sensor gives readout specific to ROCK activity

A previous ROCK sensor cross-reacted with PKA, reducing its utility in studying ROCK activity specifically^53^. To validate that RhoKAR readout is insensitive to PKA activity, 10 µM forskolin (FSK), a PKA activator was used to stimulate cells (**Figure 4A**). FSK stimulation did not significantly change RhoKAR activity compared to vehicle control (**Figure 4B**), indicating that RhoKAR is insensitive to PKA activity (**Figure 4C**). In serum-starved cells, we next inhibited PKA using 150 nM H89 and then stimulated cells with LPA during time-course imaging (**Figure 4D**). We did not observe significant differences in fluorescence anisotropies between the H89 and DMSO pretreated cells after LPA stimulation (**Figure 4E**, **4F**), indicating that sensor activity is also insensitive to PKA inhibition.

**Figure 4:**
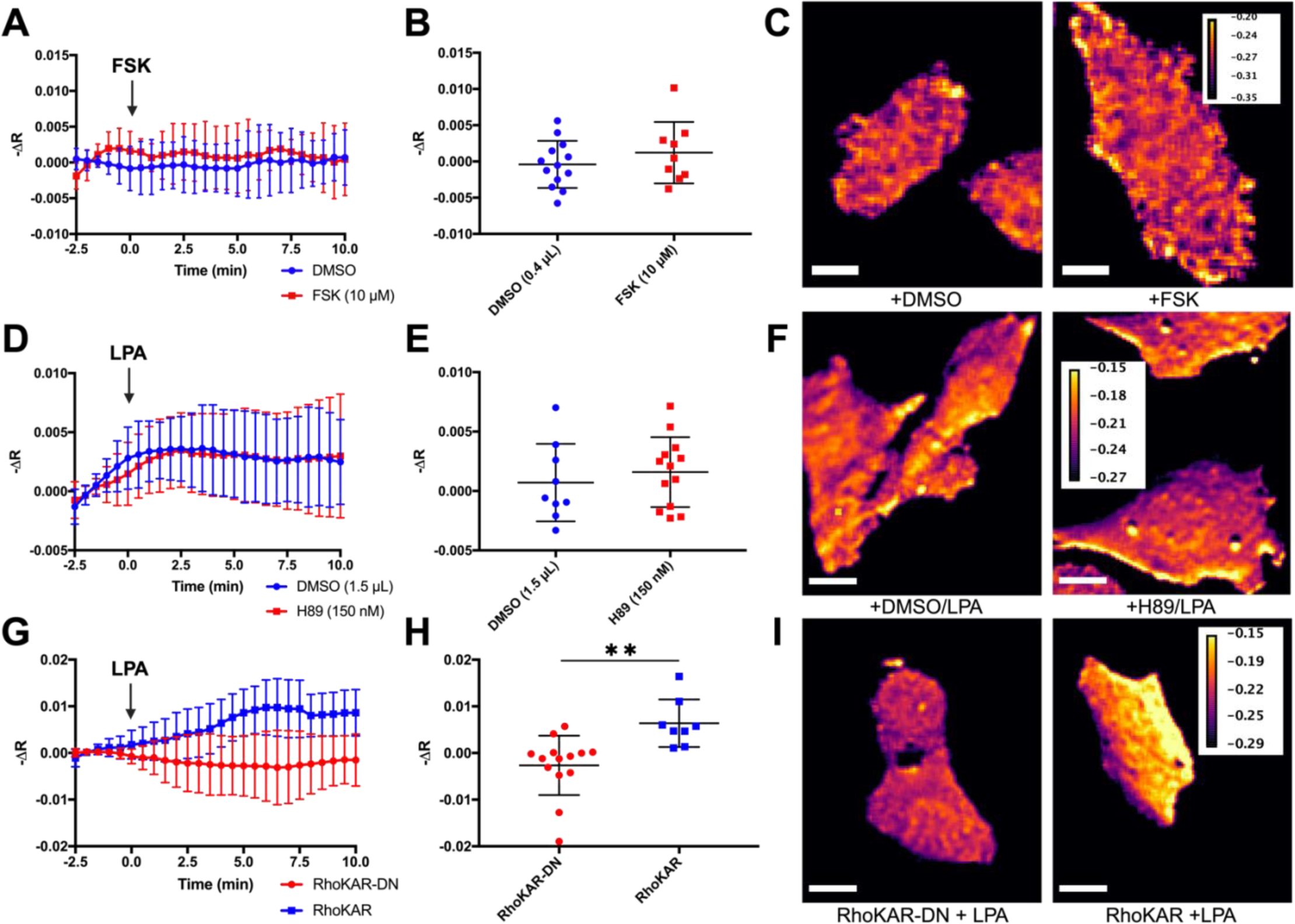
RhoKAR sensor readout is specific to ROCK activity. **A**, **B**: Time-course change in fluorescence anisotropies in individual NIH-3T3 cells expressing the RhoKAR sensor when stimulated with Forskolin or DMSO vehicle control (t-test, ns; n = 13 for DMSO, n = 9 for FSK). **C**: Representative FRET map images of cells 5 minutes after treatment with DMSO or FSK. Scale bar = 10 µm. **D**, **E**: Time-course change in fluorescence anisotropies in individual serum-starved cells expressing the RhoKAR sensor when pretreated with 150 nM H89 or 1.5 µL DMSO vehicle control and stimulated with LPA (t-test, ns; n = 9 for DMSO, n = 13 for H89). **F**: Representative FRET map images of cells pretreated with DMSO or H89 5 minutes after LPA stimulation. Scale bar = 10 µm. **G**, **H**: Time-course change in fluorescence anisotropies in individual serum-starved cells expressing the RhoKAR wild-type sensor (blue) or the RhoKAR-DN phospho-dead mutant sensor after stimulation with 6 µM LPA (t-test, **p<0.01; n = 14 for RhoKAR-DN, n = 8 for RhoKAR). **I**: Representative FRET map images of cells expressing the RhoKAR-DN or RhoKAR sensor 5 minutes after LPA stimulation. Scale bar = 10 µm.

Next, we generated a non-phosphorylatable mutant version of the sensor, the RhoKAR-DN sensor, with a threonine-to-alanine substitution at the phosphorylation site. Cells expressing the mutant or wild-type sensor were stimulated with 6 µM LPA after serum starvation (**Figure 4G**). While the wild-type RhoKAR sensor responded to LPA stimulation, the RhoKAR-DN sensor was insensitive to stimulation with 6 µM LPA, with significantly lower change in time-course fluorescence anisotropies in individual cells after LPA stimulation compared to the wild-type sensor (**Figure 4H**, **4I**). Together, these data show that RhoKAR specifically reports ROCK activity, and that the shifts we observe in anisotropies correspond to conformational change following phosphorylation of ROCK substrate on the biosensor.

### RhoKAR visualizes subcellularly compartmentalized ROCK activity

Spatial compartmentalization is an integral part of cellular processes including GPCR engagement, endocytosis, signaling for PKA, cAMP, and calcium^58–62^. Imaging techniques allowing for visualization of these dynamics has been utilized to study cyclic nucleotide signaling, small GTPase activity, as well as membrane and cytoskeletal dynamics^63–66^. RhoA activity is tightly regulated with subcellularly localized activation during cleavage furrow formation in cytokinesis, and during cell adhesion, endothelial barrier remodeling, and cell migration^15,67–70^. Whether ROCK follows the same activation patterns is unknown. To our knowledge, no biosensor exists that allows for quantitating ROCK activity specifically with high spatiotemporal resolution without crosstalk from other kinases. To visualize the dynamics of ROCK activity at a subcellular level, we imaged fibroblasts expressing the RhoKAR sensor over a one-hour time course (**Figure 5A**). Compartmentalized activation of ROCK was apparent, with areas of high ROCK activity observed during membrane protrusion and retraction over the imaging time course (**Figure 5B**). These data show that the RhoKAR sensor can visualize ROCK activity at the subcellular level.

**Figure 5:**
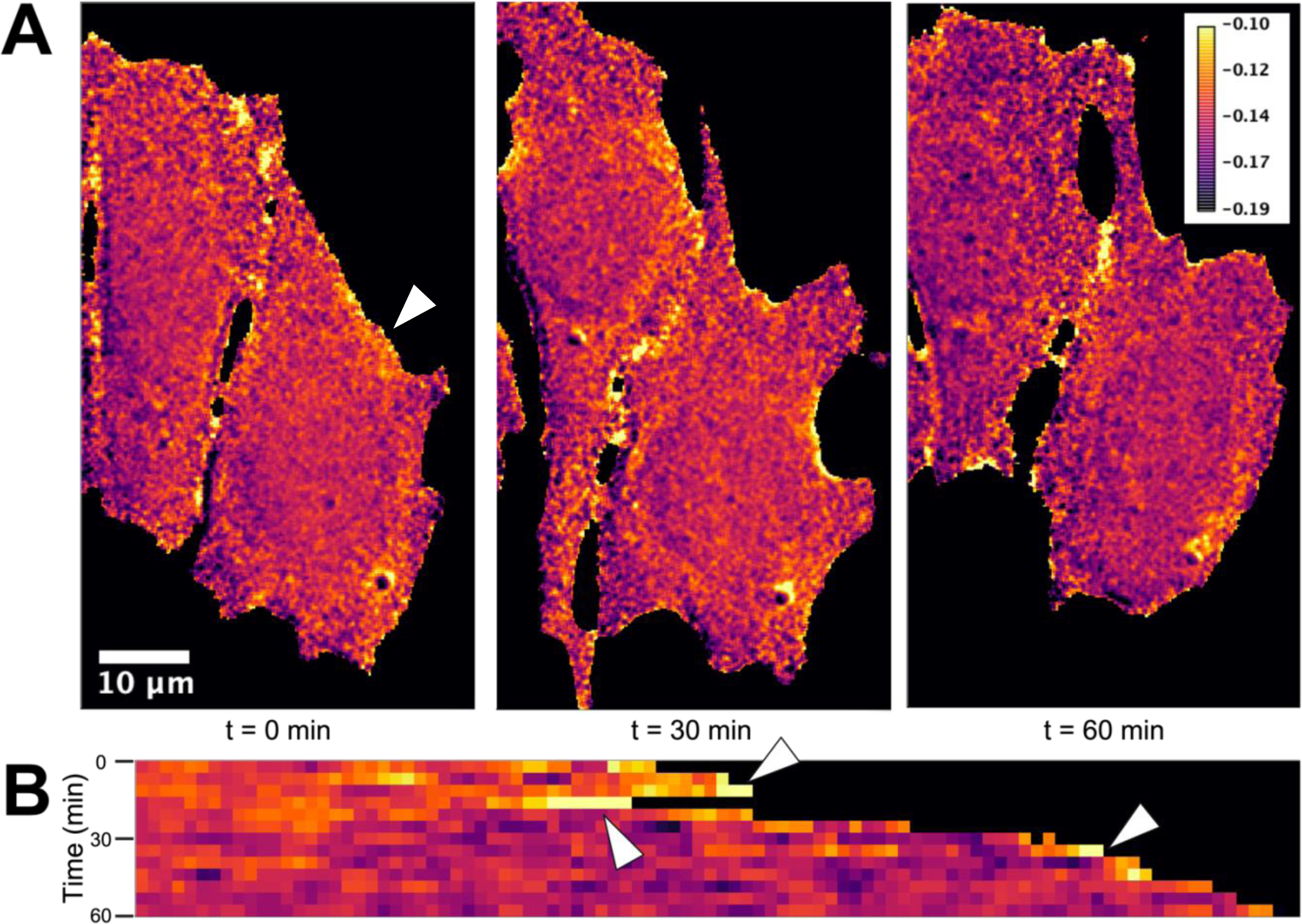
RhoKAR sensor allows for visualization of subcellular compartmentalized activation of ROCK. **A**: NIH-3T3 cells expressing the RhoKAR sensor were imaged at 40X with one image taken every 5 minutes for 1 hour. Ratio images were generated and pseudo-colored to show areas of high RhoKAR activity. During the time course, one area of the cell membrane, indicated by an arrow, protruded and retracted. **B**: Kymograph of the cell membrane protruding and retracting during course of imaging, taken at the approximate position of the arrow in the leftmost panel in A. Areas of high RhoKAR activity in the kymograph activity are demarked with arrows. High ROCK activity is observed during the process of cell protrusion and retraction.

### Calcium activates RhoA and ROCK in fibroblasts

Calcium (Ca^2+^) is a ubiquitous second messenger in eukaryotic cells^71–74^. Due to the long-observed mechanism of calcium sensitization of the contractile apparatus in vascular smooth muscle, which is mediated by active RhoA and ROCK’s inhibition of myosin light chain phosphatase^75^, and other recent reports of calcium-dependent activation of RhoA^76–84^, we were interested in exploring whether Ca^2+^ activated RhoA and ROCK in fibroblasts.

We applied ionomycin calcium salt (IM), a calcium ionophore, to cells expressing our RhoA-mCer3 two-color FRET sensor^48^ (**Figure 6A**) or the RhoKAR sensor (**Figure 6D**). Ionomycin has been used extensively in the literature to increase intracellular calcium in live cells, including in fibroblasts^71,85,86^. Both the RhoA-mCer3 (**Figure 6B**, **6C**) and RhoKAR (**Figure 6E**, **6F**) sensors showed significant increases in FRET after 2.5 µM IM treatment compared to DMSO vehicle control, indicating that Ca^2+^ stimulation activates both RhoA and ROCK in fibroblasts. Both positive and negative regulation by calcium is observed in hormone secretion and store-operated calcium channel conductivity, among other processes^87–89^. To see whether ROCK activity also shifted in response to decreasing Ca^2+^, we treated cells with 100 µM EGTA to reduce Ca^2+^ availability (**Figure 6G**). EGTA treatment significantly decreased RhoKAR activity compared to dH_2_O vehicle control (**Figure 6H**, **6I**), illustrating that reduction of intracellular calcium causes decreases in ROCK activity. Together, these results indicate that RhoA and ROCK activity in NIH-3T3 fibroblasts is calcium-dependent.

**Figure 6:**
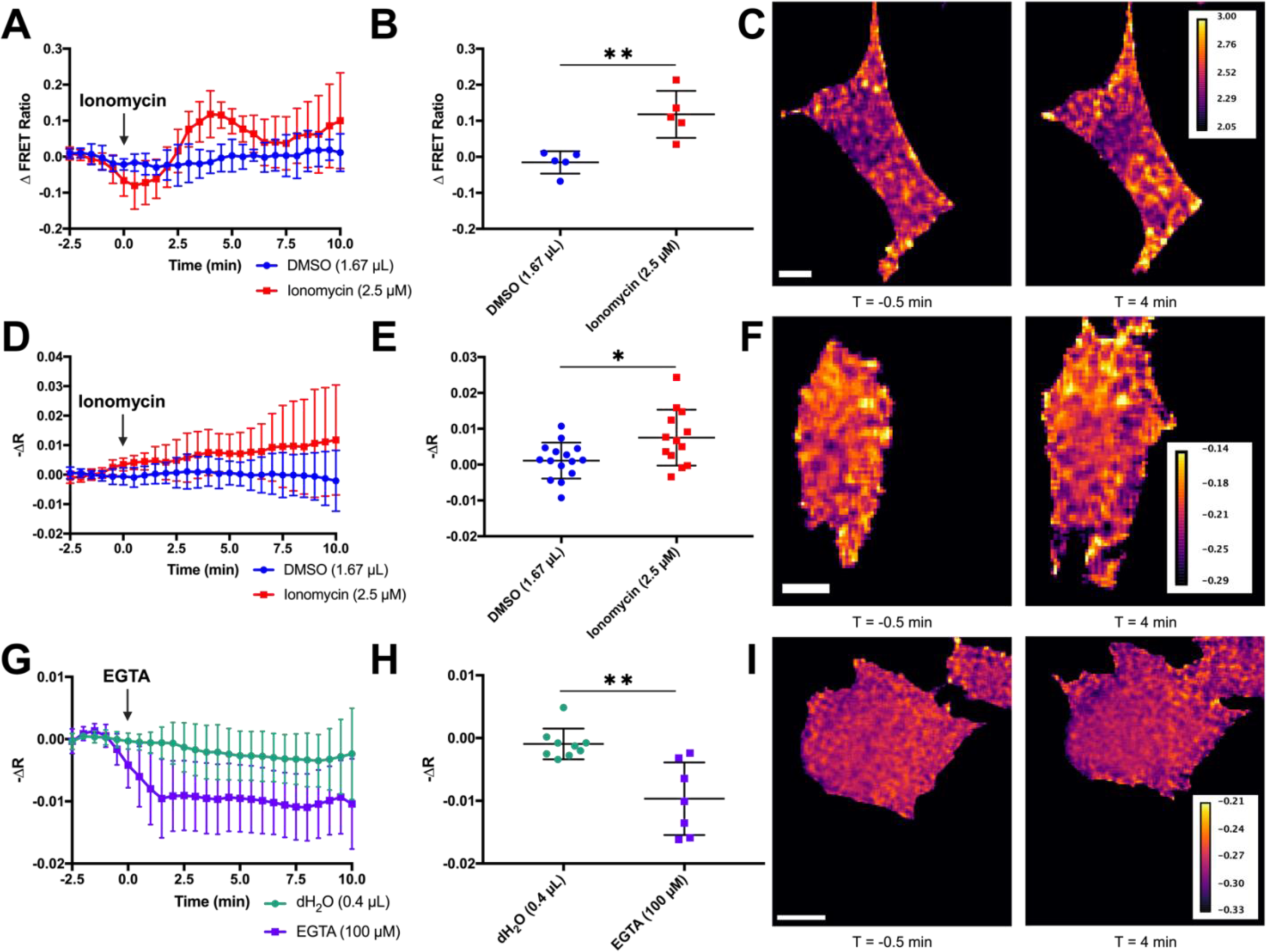
Calcium activates RhoA and ROCK in NIH-3T3 fibroblasts. **A**, **B**: Time course change in FRET in single cells expressing the RhoA two-color FRET sensor shows a significant increase in RhoA activity after ionomycin stimulation compared to DMSO vehicle control (n = 5 per treatment group, t-test **p<0.01). **C**: Representative FRET map images of cells 4 minutes after treatment with ionomycin. Scale bar = 10 µm. **D**, **E**: Time course change in fluorescence anisotropies were calculated for individual cells expressing the RhoKAR sensor after stimulation with ionomycin or DMSO vehicle control. (n = 15 for DMSO, n = 13 for ionomycin, t-test *p<0.05). **F**: Representative FRET map images of cells 4 minutes after treatment with ionomycin. Scale bar = 10 µm. **G**, **H**: Time course change in fluorescence anisotropy in individual cells expressing the RhoKAR sensor after EGTA or dH_2_O vehicle control treatment (n = 9 for dH_2_O, n = 7 for EGTA, **p<0.01). **I**: Representative FRET map images of cells 4 minutes after treatment with EGTA. Scale bar = 10 µm.

### Calcium activation of ROCK in fibroblasts requires Calmodulin/CaMKII

Following our observation of calcium-dependent activation of RhoA and ROCK in mouse fibroblasts, we wanted to determine the underlying signaling mechanism. Calmodulin (CaM) is a ubiquitously expressed and highly conserved protein that undergoes a conformational change when calcium binds to any of its four binding sites^90–92^. Active CaM in complex with Ca^2+^ can relieve the autoinhibition of Calmodulin-dependent kinase II (CaMKII), a serine-threonine protein kinase and major effector of CaM^92,93^. CaM and CaMKII drive many physiological processes, including cytoskeletal rearrangement, apoptosis, and cell proliferation^91,94,95^. Another group has reported that ionomycin stimulation activates RhoA in rabbit aortic vascular smooth muscle and that this stimulation was sensitive to the inhibition of calmodulin but not CaMKII^96^. Others have observed that CaMKII inhibition blocks RhoA activation following glutaminergic calcium influx in dendritic spines^79^. To probe the role of CaM in Ca^2+^-dependent activation of ROCK in fibroblasts, we pretreated cells expressing the RhoKAR sensor with W-7, a CaM inhibitor, or DMSO vehicle control, then stimulated with 2.5 µM ionomycin (**Figure 7A**). W-7 pretreatment significantly decreased RhoKAR response to ionomycin stimulation compared to DMSO vehicle control pretreatment (**Figure 7B**, **7C**), indicating that ionomycin-stimulated activation of ROCK requires calmodulin. Next, we chose to look downstream at CaMKII to determine whether calcium-dependent activation of ROCK in mouse fibroblasts requires this effector. KN93, a CaMKII inhibitor, was applied before stimulation with 2.5 µM ionomycin in cells expressing the RhoKAR sensor (**Figure 7D**). Compared to a DMSO vehicle control, KN93 pretreatment prevented ROCK activation by ionomycin (**Figure 7E**, **7F**), indicating that CaMKII is involved in RhoA/ROCK activation downstream of ionomycin stimulation and CaM activity in mouse fibroblasts.

**Figure 7:**
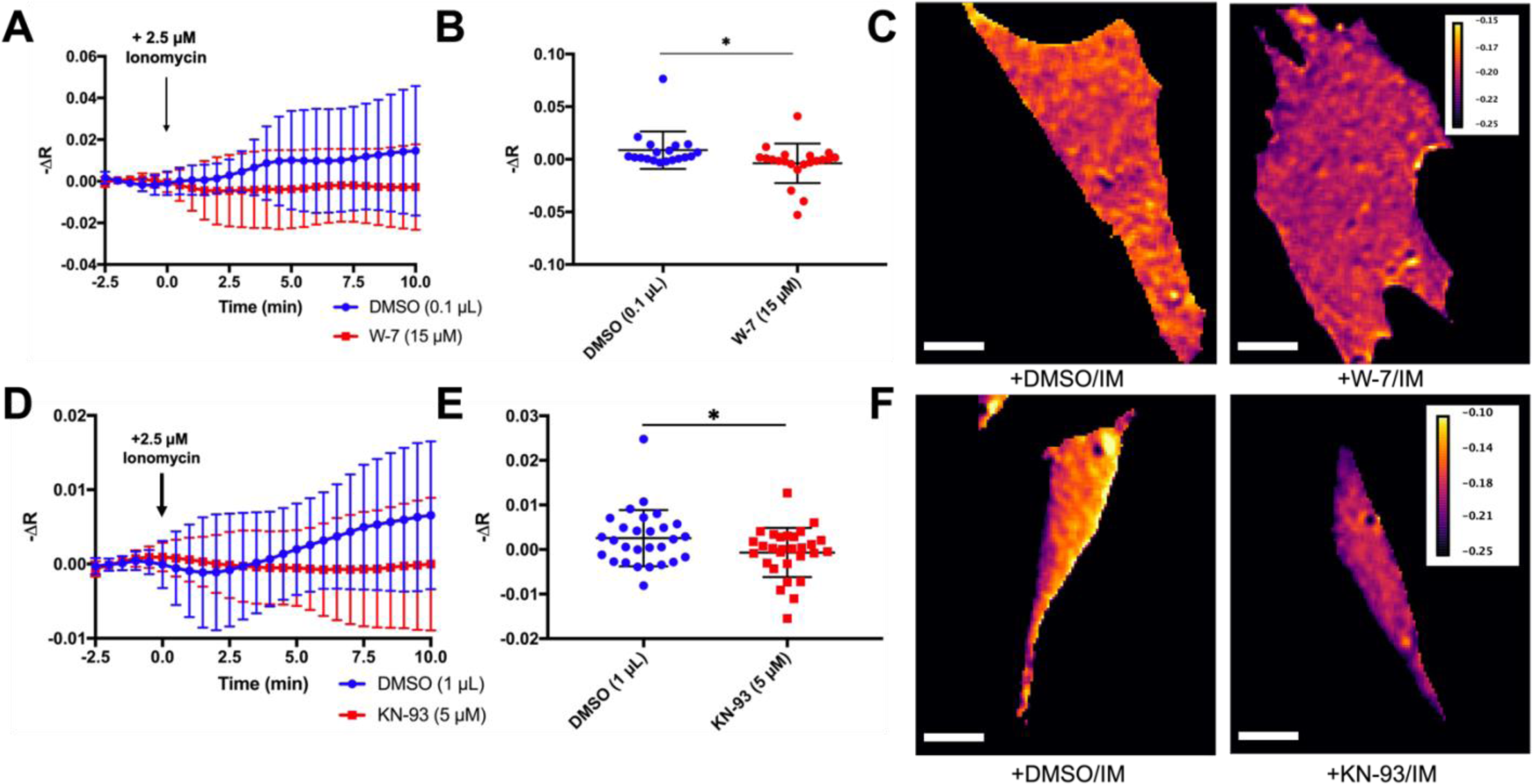
Calcium-dependent activation of ROCK requires Calmodulin and CaMKII. **A**, **B**: Time course change in fluorescence anisotropies were calculated in individual cells expressing the RhoKAR sensor after pretreatment with DMSO or W-7 and stimulation with ionomycin at T = 0 min (n = 19 for DMSO/IM, n = 21 for W-7/IM, t-test *p<0.05). **C**: Representative FRET map images of cells pretreated with DMSO or W-7 4 minutes after stimulation with ionomycin. Scale bar = 10 µm. **D**, **E**: Time course change in fluorescence anisotropies were calculated in individual cells expressing the RhoKAR sensor after pretreatment with DMSO or KN-93 and stimulation with ionomycin at T = 0 min (n = 28 for DMSO/IM, n = 29 for KN-93/IM, t-test *p<0.05). **F**: Representative FRET map images of cells pretreated with DMSO or KN-93 5.5 minutes after stimulation with ionomycin. Scale bar = 10 µm.

## DISCUSSION

### RhoKAR sensor allows for sensitive detection of changes in ROCK activity in live cells

FRET-based kinase activity reporter (KAR) type sensors have been developed for various kinases, including AKAR for PKA, CKAR for PKC, and EKAR for ERK^97–99^. These sensors have proven highly useful in studying cell signaling dynamics. Using an approach similar to that used to develop other KAR-type sensors, as well as a previously published two-color ROCK-FRET sensor^53^, we have designed a novel single-color FRET sensor for Rho-kinase (**Figure 2A, 2B**). The conformational change in the RhoKAR sensor associated with the activation of ROCK and phosphorylation of the substrate leads to a decrease in anisotropy, corresponding to an increase in FRET. Because of the higher sensitivity of anisotropy-based measurements and the large spectral occupancy of two-color FRET sensors, the single-color FRET approach is well-suited for measuring changes in dynamic cellular signaling systems such as the RhoA/ROCK pathway where it would be advantageous to be able to measure the activity of multiple proteins at once.

Stimulation cells expressing the RhoKAR sensor with classical RhoA activator LPA decreased anisotropy, indicating increased FRET and thus increased ROCK activity (**Figure 3A-3C**). Treatment with ROCK inhibitor Y27632 decreased RhoKAR activity (**Figure 3D-3F**). Because a previous ROCK-FRET sensor showed cross-reactivity to PKA^53^, we wanted to validate that RhoKAR activation is insensitive to PKA activity. PKA activation using FSK did cause significant changes to RhoKAR activity compared to vehicle control (**Figure 4A-4C**). Similarly, pretreatment of cells with H89, a PKA inhibitor, did not impact RhoKAR’s response to LPA stimulation (**Figure 4D-4F**). A non-phosphorylatable mutant of the RhoKAR sensor, RhoKAR-DN, with the phosphorylation site on the ROCK substrate mutated from threonine to alanine, showed a significantly lower response to LPA stimulation compared to the wild-type RhoKAR sensor (**Figure 4G-4I**). Together, these results indicate that our sensor gives readout specific to Rho-kinase and is insensitive to activation or de-activation of PKA, as well as that the changes in anisotropy reported by the RhoKAR sensor are due to changes in sensor conformation in response to phosphorylation of the substrate by active ROCK.

### RhoKAR sensor reveals compartmentalized activation of ROCK during cellular protrusion and retraction

RhoA must be precisely spatiotemporally regulated for a variety of cellular processes to proceed effectively. During migration, RhoA is active at both the leading edge and the retracting tail edge^8,100^. RhoA is active at the furrow in the equatorial zone but inhibited in the polar regions during cytokinesis^68,69^. Additionally, RhoA helps strengthen cell-cell interaction during adhesion after initial cell-cell contact is established^69^. Whether RhoA’s effector ROCK shares these activation patterns is not well understood.

In cells expressing the RhoKAR sensor, time course images at 40X overlaid with a FRET map reveal localized areas of high ROCK activity during cellular retraction and protrusion (**Figure 5A-B**). These areas of high RhoKAR activity illustrate the role of ROCK in cellular protrusion and retraction. RhoA’s role in driving changes to cell shape, mainly by modulating the activity and stability of the cytoskeleton, has been of interest for decades^101^. Because of RhoA/ROCK’s well-characterized role in promoting the active state of the actomyosin contractile apparatus via inhibitory phosphorylation of myosin light chain kinase ^75,102,103^, this localized increase in ROCK activity may be a significant driving force in the cytoskeletal processes pushing forward or pulling back the membrane.

### Calcium activates RhoA and ROCK in fibroblasts via calmodulin/CaMKII signaling

Calcium is a vital second messenger in the cell^71–74^. Resting Ca^2+^ concentrations in the cytoplasm are in the nanomolar range, while the ER and extracellular space maintain Ca^2+^ concentrations in the millimolar range. Rapid influxes of Ca^2+^ to the cytoplasm or microdomains within the cell is used to drive fast processes and responses to signals such as GPCR engagement or cell depolarization^104–106^. Like RhoA/ROCK activity, calcium must be precisely spatiotemporally regulated for physiological processes, such as proliferation and migration, to proceed effectively^61,107,108^. Recent findings in the literature have shown Ca^2+^-dependent activation of RhoA in a variety of tissue types, including vascular, sphincteric, and airway smooth muscle, as well as hippocampal neurons, cerebellar granule cells, embryonic kidney cells, endothelial cells, and intestinal epithelial cells^77–81,109^. Calcium also activates RhoA in disease states, including brain, colon, and endometrial cancer^82–84^.

To determine whether calcium activates RhoA/ROCK in fibroblasts, cells expressing our RhoA-mCer3 two-color FRET sensor or the RhoKAR sensor were treated with ionomycin, a calcium ionophore. Following ionomycin treatment, we observed increases in both RhoA and ROCK activity (**Figure 6A-F**). Decreasing calcium with EGTA also decreased ROCK activity (**Figure 6G-I**). These data indicate that both RhoA and ROCK activation are calcium-dependent in fibroblasts. Ionomycin stimulation also activates RhoA in rabbit aortic vascular smooth muscle. In this system, activation of RhoA by ionomycin was sensitive to treatment with W-7, a calmodulin inhibitor, but not KN93, a CaMKII inhibitor^96^. However, another group has reported that in the dendritic spines of rat hippocampal neurons, CaMKII inhibition was sufficient to prevent RhoA activation following glutaminergic signaling dependent calcium influx^79^. Similarly, we observed that ROCK activation by ionomycin was sensitive to W-7 pretreatment (**Figure 7A-C**). CaMKII inhibition with KN-93 in fibroblasts also prevented ionomycin-dependent activation of ROCK (**Figure 7D-F**). The KN-93 sensitivity observed in mouse fibroblasts and rat hippocampal neurons, but not rabbit vascular smooth muscle, suggests that mechanisms for calcium-dependent activation are tissue-specific.

Taken together, our findings show that an ionomycin-dependent increase in intracellular calcium activates RhoA and ROCK via CaM and CaMKII. Although characterizing any GEFs potentially involved in the proposed signaling pathway was beyond the scope of this work, several GEFs activate downstream of CaM and CaMKII, including ARHGEF23, ARHGEF24, and ARHGEF39^110,111^. Other groups have observed transient, localized increases and decreases in RhoA activity due to changes in intracellular calcium during epithelial tight junction remodeling^112^. Our findings in fibroblasts have implications for processes such as cell migration, where an influx of calcium could promote RhoA/ROCK activity and the creation of contractile forces via cytoskeletal activation or reorganization.

## Biosensor Availability

Plasmids for biosensors developed for this paper will be made publicly available via Addgene.

## Author Contributions

A.E.M. collected and analyzed the data, generated figures, and wrote the manuscript. M.A.R. conceived the experiments and edited the manuscript.

